# Hypoxic response regulators RHY-1 and EGL-9/PHD promote longevity through a VHL-1 independent transcriptional response

**DOI:** 10.1101/2020.03.12.989061

**Authors:** Joseph Kruempel, Hillary A. Miller, Megan L. Schaller, Abrielle Fretz, Marshall Howington, Marjana Sarker, Shijiao Huang, Scott F. Leiser

**Affiliations:** Molecular & Integrative Physiology Department, University of Michigan, Ann Arbor, MI 48109, USA; Cellular and Molecular Biology Graduate Program, University of Michigan, Ann Arbor, MI 48109, USA; Department of Internal Medicine, University of Michigan, Ann Arbor, MI 48109, USA

**Keywords:** *C. elegans*, lifespan, aging, HIF-1 signaling, hypoxic response, RHY-1, EGL-9/PHD

## Abstract

HIF-1-mediated adaptation to changes in oxygen availability is a critical aspect of healthy physiology. HIF is regulated by a conserved mechanism whereby EGLN/PHD family members hydroxylate HIF in an oxygen-dependent manner, targeting it for ubiquitination by Von-Hippel-Lindau (VHL) family members, leading to its proteasomal degradation. The activity of the only *C. elegans* PHD family member, EGL-9, is also regulated by a hydrogen sulfide sensing cysteine-synthetase-like protein, CYSL-1, which is, in turn, regulated by RHY-1/acyltransferase. Over the last decade multiple seminal studies have established a role for the hypoxic response in regulating longevity, with mutations in *vhl-1* substantially extending *C. elegans* lifespan through a HIF-1-dependent mechanism. However, studies on other components of the hypoxic signaling pathway that similarly stabilize HIF-1 have shown more mixed results, suggesting that mutations in *egl-9* and *rhy-1* frequently fail to extend lifespan. Here, we show that *egl-9* and *rhy-1* mutants suppress the long-lived phenotype of *vhl-1* mutants. We also show that RNAi of *rhy-1* extends lifespan of wild-type worms while decreasing lifespan of *vhl-1* mutant worms. We further identify VHL-1-independent gene expression changes mediated by EGL-9 and RHY-1 and find that a subset of these genes contributes to longevity regulation. The resulting data suggest that changes in HIF-1 activity derived by interactions with EGL-9 likely contribute greatly to its role in regulation of longevity.

## Introduction

Adaptation to changes in oxygen availability is a central requirement for aerobic life. In response to hypoxia, reduced oxygen-dependent hydroxylation of Hypoxia Inducible Factor α (HIFα) transcription factors by members of the EGLN/Proline-Hydroxylase (PHD) family triggers stabilization of HIFα proteins and activation of a transcriptional stress response that promotes survival (Epstein et al., 2001). This hypoxic response plays critical roles in a variety of pathological conditions including inflammation and cancer (Balamurugan, 2016; Pezzuto & Carico, 2018; Ramakrishnan & Shah, 2016). Constitutive stabilization of the sole *C. elegans* HIFα family member, HIF-1, by deletion of the Von-Hippel-Lindau ubiquitin ligase, VHL-1, which ubiquitinates HIF-1 and targets it for degradation, results in HIF-1-dependent increases in stress response and longevity (Cockman et al., 2000; Jiang et al., 2001; Leiser & Kaeberlein, 2010; Mehta et al., 2009; Müller et al., 2009; Treinin et al., 2003; Zhang et al., 2009).

Genetic studies in *C. elegans* have identified additional players in the hypoxic signaling pathway. Activity of EGL-9, the only known *C. elegans* PHD family member, is inhibited by direct interaction with the H_2_S sensing cysteine-synthetase family member CYSL-1 (Ma et al., 2012). CYSL-1 protein levels are in turn reduced through an unknown mechanism by Regulator of Hypoxia-inducible factor-1 (RHY-1), an ER transmembrane protein with predicted acyltransferase activity (Ma et al., 2012; Shen et al., 2006). Predicted loss-of-function mutations in *rhy-1* stabilize HIF-1 and produce expression patterns of HIF-1 target genes that are consistent with reduced EGL-9 activity (Shen et al., 2006).

Interestingly, while *vhl-1* mutation extends *C. elegans* lifespan across culture conditions, the role of EGL-9/PHD is more context dependent. While *egl-9(RNAi)* extends lifespan at 20°C, the lifespan phenotypes of partial loss-of-function mutations in *egl-9* are temperature-dependent, extending lifespan in a HIF-1-dependent manner at low temperatures (15°C) but not at higher temperatures (20°C and 25°C) (Leiser et al., 2011; Leiser & Kaeberlein, 2010; Miller et al., 2017; Zhang et al., 2009). The loss-of-function mutant *egl-9(sa307)* also reduces the lifespan of dietary restricted animals and long-lived *rsks-1* mutants when animals are cultured at 25°C, suggesting that EGL-9 activity may promote lifespan in multiple contexts (Di Chen & Kapahi, 2009). Furthermore, recent work from our lab showed that *rhy-1* putative knockout mutants were not long-lived at any temperature, despite their reported robust activation of hypoxic response genes (Miller et al., 2017; Shen et al., 2006).

Previous studies on the roles of EGL-9, RHY-1, and VHL-1 show that 1) HIF-1 stabilization when *vhl-1* is mutated leads to robust induction of *egl-9* and *rhy-1*, 2) EGL-9 and RHY-1 have VHL-1-independent effects on transcription of some hypoxic response genes, and 3) EGL-9 and RHY-1 play a VHL-1-independent role in pathogen and hydrogen sulfide resistance (Horsman et al., 2019; Luhachack et al., 2012; Shao et al., 2009; Shao et al., 2010; Shen et al., 2005; Shen et al., 2006). However, the possibility that EGL-9 and RHY-1 modulate longevity through a downstream, VHL-1-independent transcriptional response has not been addressed. Here we present a genetic study demonstrating that EGL-9 and RHY-1 are necessary for lifespan extension when HIF-1 is stabilized by *vhl-1* mutation. We show that, like EGL-9, RHY-1 has both longevity promoting and inhibiting activities. We further identify genes that are oppositely regulated in *vhl-1* and *egl-9* or *rhy-1* mutants, suggesting that RHY-1 and EGL-9 promote a VHL-1-independent transcriptional response when HIF-1 is stabilized by *vhl-1* mutation. Lastly, we find that RNAi knockdown of four genes downregulated in *vhl-1(ok161)* mutants and upregulated in *egl-9(sa307)* mutants, with likely functions in innate immunity, each partially rescues lifespan extension in *egl-9(sa307);vhl-1(ok161)* mutants. Together, our results suggest that EGL-9 modulates lifespan by regulating a VHL-1-independent transcriptional program.

## Results

### RHY-1 and EGL-9 promote longevity in *vhl-1* mutants

We initially hypothesized that the presence of EGL-9 and RHY-1 is necessary to promote longevity downstream of HIF-1 stabilization. This hypothesis would suggest that reduced activity of EGL-9 or RHY-1 would prevent or mitigate lifespan extension when HIF-1 is stabilized by reducing VHL-1 activity. Previous studies show that *vhl-1* and *egl-9* mutant strains show similar lifespan phenotypes at low temperatures (15°C), making epistasis experiments difficult to interpret (Miller et al., 2017). However, at high temperatures, *egl-9(sa307)* mutants have short to wild-type lifespans while *vhl-1(ok161)* mutants are long-lived (Di Chen & Kapahi, 2009; Leiser et al., 2011; Miller et al., 2017). Thus, we tested whether *egl-9(sa307)* and/or *rhy-1(ok1402)* mutants suppress the extended longevity phenotype of *vhl-1(RNAi)*-treated or *vhl-1(ok161)* mutant animals at high temperature (25°C). Our results (**Figure 1**) show that *rhy-1(ok1402)* **(Fig. 1A)** and *egl-9(sa307)* **(Fig. 1B)** mutants abrogate the extended longevity phenotype caused by *vhl-1(RNAi)* at 25°C. Furthermore, *rhy-1(ok1402)* fully abrogated the extended longevity phenotype of *vhl-1(ok161)* mutants **(Fig. 1C)**, while *egl-9(sa307)* mutants partially suppressed the extended longevity phenotype *vhl-1(ok161)* mutants at 25°C **(Fig. 1D)**. We also observed that *rhy-1(RNAi)* reduces the lifespan of *vhl-1* mutants at low temperatures (15°C, **(Fig. 1E)**) and that the lifespan of *rhy-1(ok1402)* mutants are not extended by *vhl-1(RNAi)* at low temperatures (15°C, **(Fig. S1)**), suggesting that these interactions are not fully temperature-dependent. These results are consistent with a model where RHY-1 and EGL-9 act in the same pathway to promote lifespan downstream of HIF-1 stabilization.

**Figure 1.**
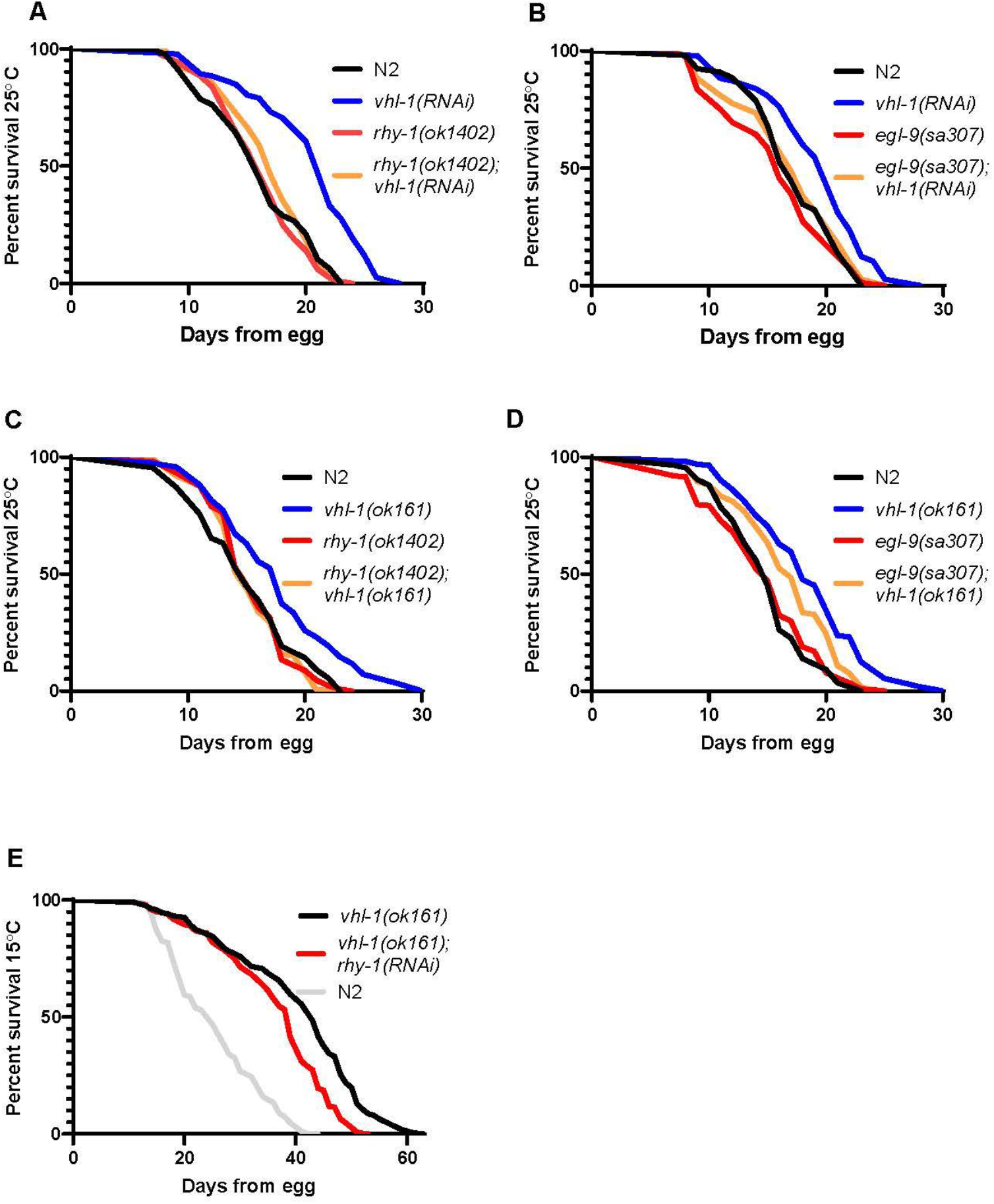
Epistasis of hypoxic response regulators. A) Lifespans of N2 (wild-type) or *rhy-1(ok1402)* mutants treated with empty vector or *vhl-1(RNAi)* at 25°C. B) Lifespans of N2 and *egl-9(sa307)* animals treated with empty vector and *vhl-1(RNAi)* at 25°C. C) Lifespans of N2, *vhl-1(ok161)*, *rhy-1(ok1402)* and *rhy-1(ok1402);vhl-1(ok161)* animals at 25°C. D) Lifespans of N2, *egl-9(sa307)*, *vhl-1(ok161)* and *egl-9(sa307);vhl-1(ok161)* animals at 25°C. E) Lifespans of vhl-1(ok161) animals treated with empty vector and *rhy-1(RNAi)* at 15°C, along with N2 controls, (p<.0001 by log-rank). Interactions of *egl-9(sa307)* and *rhy-1(ok161)* with *vhl-1(ok161)* and *vhl-1(RNAi)* were statistically different than interactions of N2 animals with *vhl-1(ok161)* and *vhl-1(RNAi)* by non-overlapping 95% confidence intervals of Mantel-Haenzel hazard ratios (**Table S1**). Data are aggregated from at least three independent experiments (**Table S2**).

### RHY-1 has longevity promoting and inhibiting activities

In the course of testing the effects of *rhy-1(RNAi)* on *vhl-1(ok161)* mutants at 15°C, we observed that *rhy-1(RNAi)* substantially extended the lifespan of wild-type control animals **(Fig. 2A)**. This result was surprising since we recently reported a study of the interaction between temperature and longevity and found that *rhy-1(ok1402)* mutants did not extend lifespan at any of the three temperatures (15°C, 20°C, or 25°C) that are commonly used for longevity studies (Miller et al., 2017). We outcrossed *rhy-1(ok1402)* five additional times to wild-type to further eliminate any background mutations and repeated lifespan experiments. Consistent with our published data, the outcrossed *rhy-1(ok1402)* strain was not long-lived: observed lifespans were modestly longer than wild-type in two of four trials, identical to wild-type in one trial, and shorter than wild-type in one trial **(Fig. 2B, S1)**. Lifespan extension by *rhy-1(RNAi)* was abrogated by *rhy-1(ok1402)*, consistent with the longevity extension phenotype caused by *rhy-1(RNAi)* being dependent on modulation of the RHY-1 gene product **(Fig. 2C)**. Furthermore, *rhy-1(RNAi)* treated *cysl-1(ok762)* and *hif-1(ia4)* mutants are short lived relative to *rhy-1(RNAi)* treated wild-type animals, consistent with a model where HIF-1 and CYSL-1 activity are required for full lifespan extension by *rhy-1(RNAi)* **(Fig. 2D-E)**. Importantly, *rhy-1(RNAi)* treatment does extend the lifespans of *cysl-1(ok762)* and *hif-1(ia4)* mutants relative to vector treated controls (**Figure 2 D,E**) suggesting that it may modulate lifespan through a mechanism that is partially independent of the CYSL-1, HIF-1 pathway. The distinct phenotypes observed in *rhy-1(ok1402)* and *rhy-1(RNAi)* are similar to the reported differences between *egl-9(RNAi)*, which extends lifespan at 20°C, and the *egl-9(sa307)* mutant, which extends lifespan at 15°C but not 20°C or 25°C (Di Chen & Kapahi, 2009; Mehta et al., 2009; Miller et al., 2017; Zhang et al., 2009). These results are consistent with a model where partial reductions in activity of *rhy-1* or *egl-9* increase longevity by stabilizing HIF-1, while stronger reduction of *rhy-1* or *egl-9* causes a secondary effect that limits lifespan downstream of HIF-1 stabilization.

**Figure 2.**
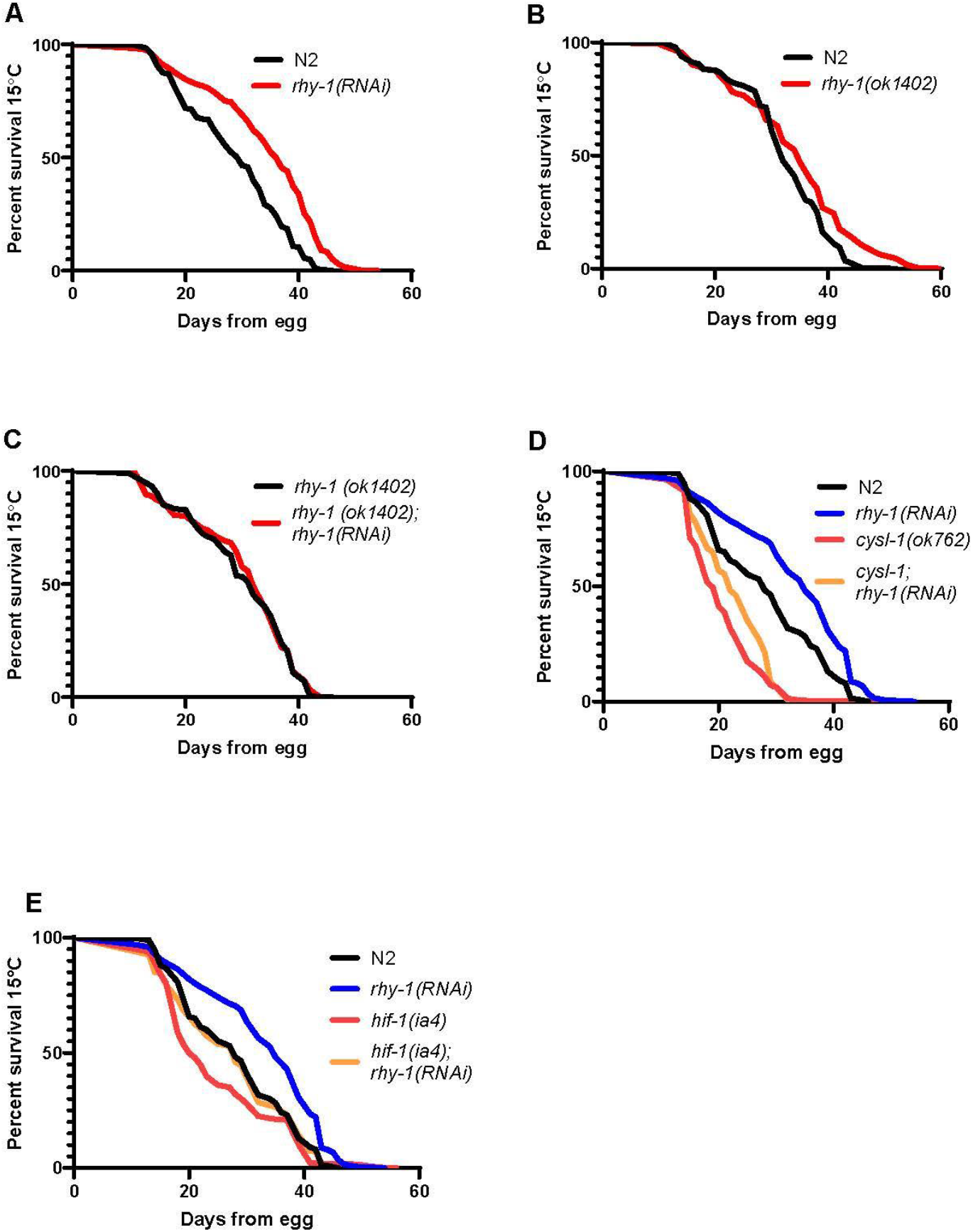
*rhy-1-* mediated regulation of longevity. A) Lifespans of N2 (wild-type) animals treated with empty vector or *rhy-1(RNAi)* (p < 0.0001 by log-rank). B) Lifespans of N2 and *rhy-1(ok1402)* animals (p < 0.0001 by log-rank). C) Lifespans of *rhy-1(ok1402)* animals treated with empty vector or *vhl-1(RNAi)*. D) Lifespans of N2 and *cysl-1(ok762)* animals treated with vector or rhy-1(RNAi) (p < 0.001 for *cysl-1(ok762);rhy-1(RNAi)* vs *rhy-1(RNAi)* by log rank) E) Lifespans of *hif-1(ia4)* and N2 animals treated with empty vector or *rhy-1(RNAi)* (p < 0.0001 for *hif-1;rhy-1(RNAi)* vs *rhy-1(RNAi)*). Interaction with *rhy-1(RNAi)* treatment was statistically different between N2 and *hif-1(ia4)* as determined by non-overlapping 95% confidence intervals of Mantel-Haenzel hazard ratios. Hazard ratios for *cysl-1(ok762);rhy-1(RNAi)*/*cysl-1(ok762)* and *rhy-1(RNAi)*/N2 were (0.58-0.82) and (0.44-0.64) respectively. Lifespan data are aggregated from at least three experiments (Table S1,S2).

### RHY-1 and EGL-9 control a VHL-1-independent transcriptional response

Previous studies reported that *egl-9* causes *vhl-1*-independent changes in expression of some transcripts (Shao et al., 2009; Shen et al., 2005; Shen et al., 2006). We hypothesized that the dominance of the *egl-9(sa307)* and *rhy-1(ok1402)* lifespan phenotypes over the *vhl-1(ok161)* lifespan phenotype might be caused by genes whose transcription is regulated by EGL-9 or RHY-1. Concurrently with our work, Angeles et al. published an analysis of transcriptome profiles from *rhy-1(ok1402)*, *egl-9(sa307)*, *hif-1(ia4)*, *egl-9;hif-1*, and *egl-9;vhl-1* mutants. They identified a class of genes whose transcription is regulated by EGL-9 in a way that is distinct from, and dominant over, their regulation by VHL-1. We will refer to this class as EGL-9/VHL-1 antagonistic genes.

To identify genes that might modulate lifespan downstream of the hypoxic response, we had previously profiled the transcriptomes of *egl-9(sa307)*, *vhl-1(ok161), rhy-1(ok402)*, and *hif-1(ia4)* mutants. We reanalyzed our data using the methodology that Angeles et al. reported and an up-to-date bioinformatic pipeline (Angeles-Albores et al., 2018; Bray et al., 2016; Pimentel et al., 2017). Our datasets identified a subset of genes that were differentially expressed in the HIF-1 negative regulator mutants relative to wild-type, further confirming that these changes are implicated in the hypoxic response, **(Fig. 3A, Table 1, Table S3,S4**). Conversely, we observed low overlap between the datasets for genes differentially expressed in the *hif-1(ia4)* background, suggesting that differences between *hif-1(ia4)* and wild-type in individual datasets may largely reflect strain-specific effects rather than HIF-1-dependent transcription under normoxia **(Fig 3B)**.

**Figure 3.**
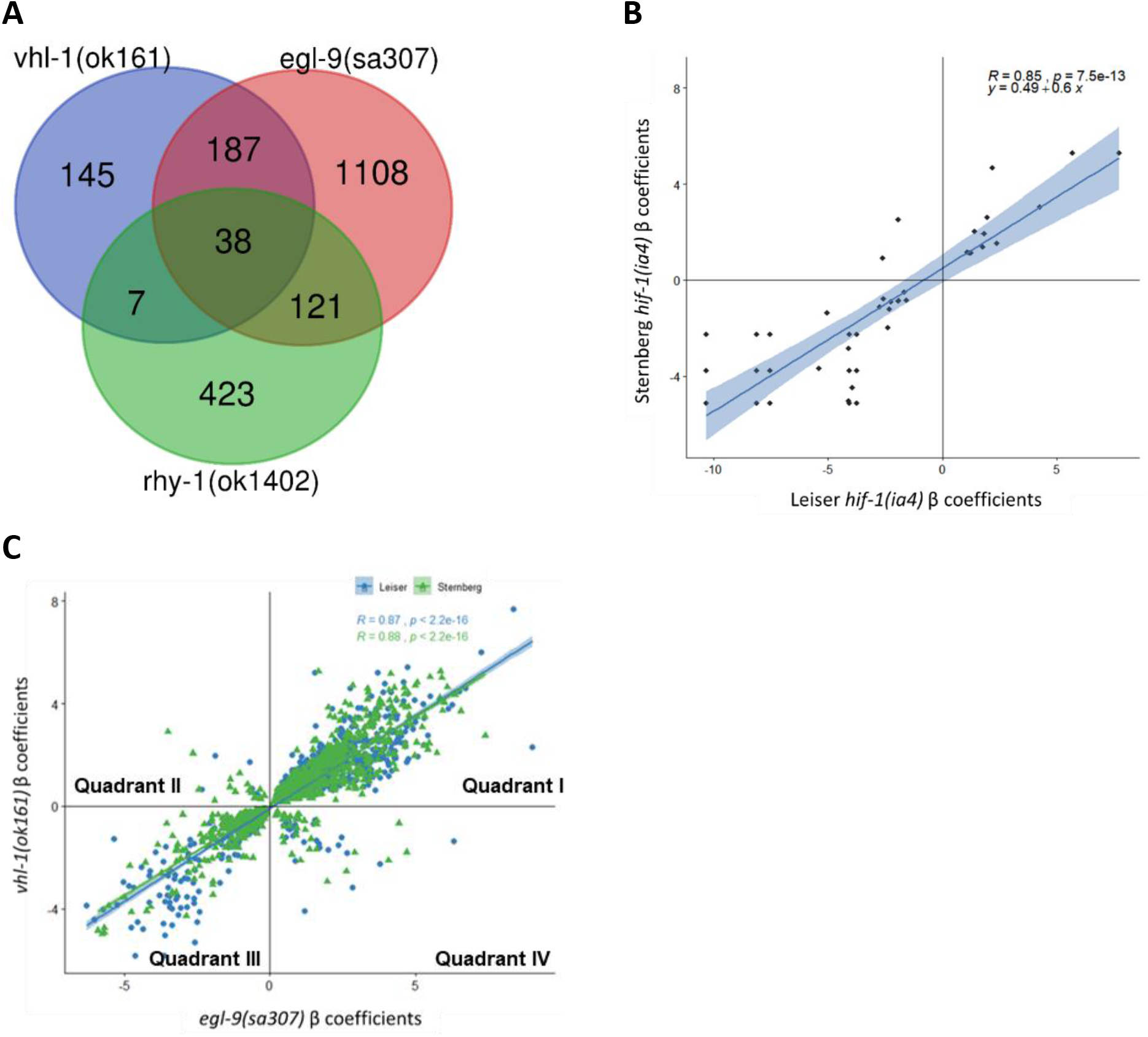
A subset of genes are antagonistically regulated by *egl-9* and *vhl-1*. A) Overlap between genes that are differentially expressed in both Leiser and Sternberg datasets for *vhl-1(ok161)* and *egl-9(sa307)* along with differentially expressed genes from Sternberg *rhy-1(ok1402)*. B) Overlap between differentially expressed genes in *hif-1(ia4)* in Leiser and Sternberg datasets. C) Correlation of expression levels of differentially expressed genes in *egl-9(sa307)* and *vhl-1(ok161)* in Leiser and Sternberg datasets.

**Table 1.** Core Hypoxic response genes. List of 38 genes that were differentially regulated in *rhy-1(ok1402)* (Sternberg only), *egl-9(sa307)* and *vhl-1(ok161)* mutants in both Leiser and Sternberg RNA-seq datasets.

| ORF | Gene | Description |
| --- | --- | --- |
| K08C7.5 | fmo-2 | FMO5 (flavin containing dimethylaniline monooxygenase 5) ortholog |
| T05B4.2 | nhr-57 | nuclear hormone receptor family transcription factor |
| C14C6.5 | C14C6.5 | predicted role in innate immune response |
| F22D6.10 | col-60 | collagen structural protein in cuticle |
| F26H9.5 | F26H9.5 | PSAT1 (phosphoserine aminotransferase 1) ortholog |
| C08E8.3 | C08E8.3 | unknown function |
| K06A4.7 | K06A4.7 | unknown function |
| F15B9.1 | far-3 | fatty acid binding protein |
| F38B2.4 | F38B2.4 | AK1 (adenylate kinase 1) ortholog |
| Y37A1B.5 | Y37A1B.5 | SELENBP1 (selenium binding protein 1) ortholog |
| F38A3.2 | ram-2 | structural protein in cuticle |
| F28F8.2 | acs-2 | ACSF2 (acyl-CoA synthetase family member 2) ortholog |
| F08G5.4 | col-130 | collagen structural protein in cuticle |
| T10D4.13 | ins-19 | predicted insulin protein |
| C31C9.2 | C31C9.2 | PHGDH (phosphoglycerate dehydrogenase) ortholog |
| F22B5.4 | F22B5.4 | unknown function |
| F10D2.9 | fat-7 | SCD5 (stearoyl-CoA desaturase 5) ortholog |
| R08E5.3 | R08E5.3 | methyltransferase orthologs |
| Y22F5A.4 | lys-1 | lysozyme activity, predicted role in innate immune response |
| C05E7.1 | C05E7.1 | unknown function |
| Y58A7A.1 | Y58A7A.1 | SLC31A1 (solute carrier family 31 member 1) ortholog |
| M05D6.6 | taap-1 | FAM162A/B (family with sequence similarity 162 member A/B) ortholog |
| W07A12.7 | rhy-1 | Negative regulator of HIF-1 |
| M199.4 | clec-190 | C-type lectin predicted to have carbohydrate binding activity |
| B0222.6 | col-144 | collagen structural protein in cuticle |
| M05D6.5 | M05D6.5 | HIGD2A/B (HIG1 hypoxia inducible domain family member 2A/B) ortholog |
| F09A5.9 | ttr-34 | predicted transthyretin-like protein |
| W07A12.6 | oac-54 | predicted amino-acyl transferase protein |
| F02H6.5 | sqrd-1 | SQOR (sulfide quinone oxidoreductase) ortholog |
| ZK637.13 | glb-1 | predicted role in heme/oxygen binding and carrying |
| F54C9.4 | col-38 | collagen structural protein |
| Y22F5A.5 | lys-2 | lysozyme activity, predicted role in innate immune response |
| F37B1.8 | gst-19 | HPGDS (hematopoietic prostaglandin D synthase) ortholog |
| T03F1.11 | T03F1.11 | predicted for CIBs (calcium and integrin binding family member) ortholog |
| VF13D12L.3 | VF13D12L.3 | predicted oxidoreductase protein |
| C04G6.2 | C04G6.2 | unknown function |
| K10B3.7 | gpd-3 | GAPDH (glyceraldehyde-3-phosphate dehydrogenase) ortholog |
| R02E4.3 | R02E4.3 | unknown function |

We next plotted β coefficients of differentially expressed genes shared between *vhl-1(ok161)* and *egl-9(sa307)* in Leiser and Sternberg datasets **(Fig 3C)**. Both datasets produce a similar pattern, with *egl-9(sa307)*, and *vhl-1(ok161)* causing highly correlated changes in expression for most shared differentially expressed genes. Both datasets also contained EGL-9/VHL-1 antagonistic genes, which were either upregulated in *vhl-1(ok161)* and downregulated in *egl-9(sa307)* (**Fig. 3C**, quadrant II), or upregulated in *egl-9(sa307)* or *rhy-1(ok1402)* and downregulated in *vhl-1(ok161)* (**Fig. 3C**, quadrant IV). The genes in quadrants II and IV, who we hypothesized could play a role in different outcomes between *vhl-1(ok161)* and *egl-9(sa307)* or *rhy-1(ok1402)* strains, are listed in **Table S5**. Together, our and the Sternberg lab’s results show that HIF-1 stabilization through loss of its negative regulators has both many common effects and a smaller number of opposing effects depending on which negative regulator is mutated.

### Knockdown of egl-9 target genes rescues lifespan *egl-9(sa307);vhl-1(ok161)* mutants

We next tested the hypothesis that EGL-9/VHL-1 antagonistic genes regulate longevity. Previous results suggest that individual longevity-pathway-target-genes often have small effects on lifespan, and longevity increases are more likely than longevity decreases to reflect modulation of the aging process as a whole (Murphy et al., 2003). Thus, we identified candidate EGL-9/VHL-1 antagonistic genes whose downregulation in *vhl-1(ok161)* mutants was reversed in *egl-9(sa307);vhl-1(ok161)* mutants and determined whether RNAi targeting them could extend life lifespan of *egl-9;vhl-1(ok161)* mutants **(Figure 4 A,C,E,G, Table S6)**.

**Figure 4.**
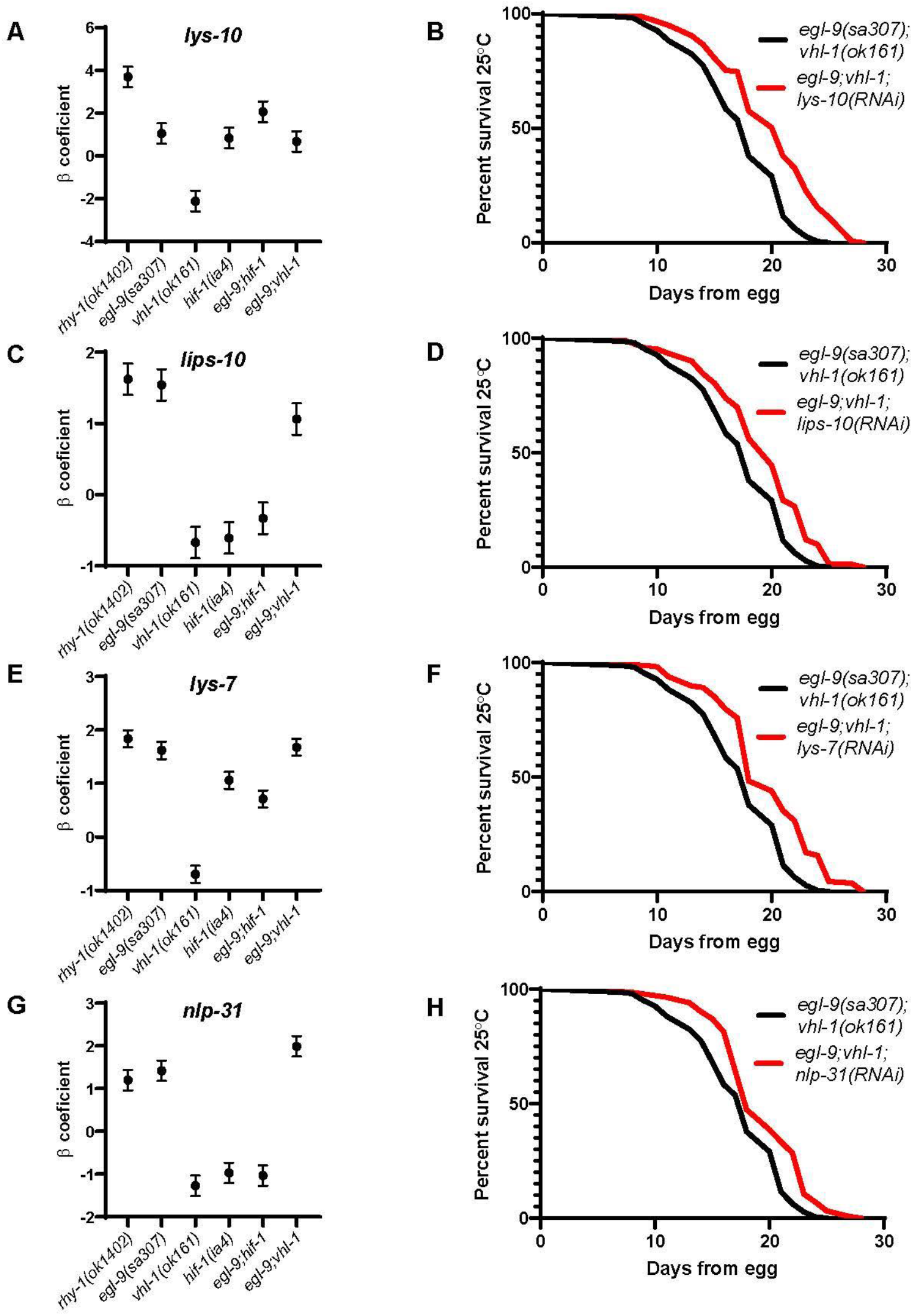
VHL-1/EGL-9 antagonistic HIF-1 targets rescue lifespan in *egl-9;vhl-1* mutants. A, C, E, G) expression levels of selected transcripts from RNA-seq analyses. B-D-F-H) Treatment with *nlp-31(RNAi)*, *lys-7(RNAi)*, *lys-10(RNAi)* and *lips-10(RNAi)* increases lifespan of *egl-9(sa307);vhl-1(ok161)* mutants (p<.05 by log-rank). Lifespan data are aggregated from at least three experiments, and are significant (p<.05 by log-rank with Bonferroni correction) in at least 3 of 5 individual trials (Table S2).

**Figure 5.**
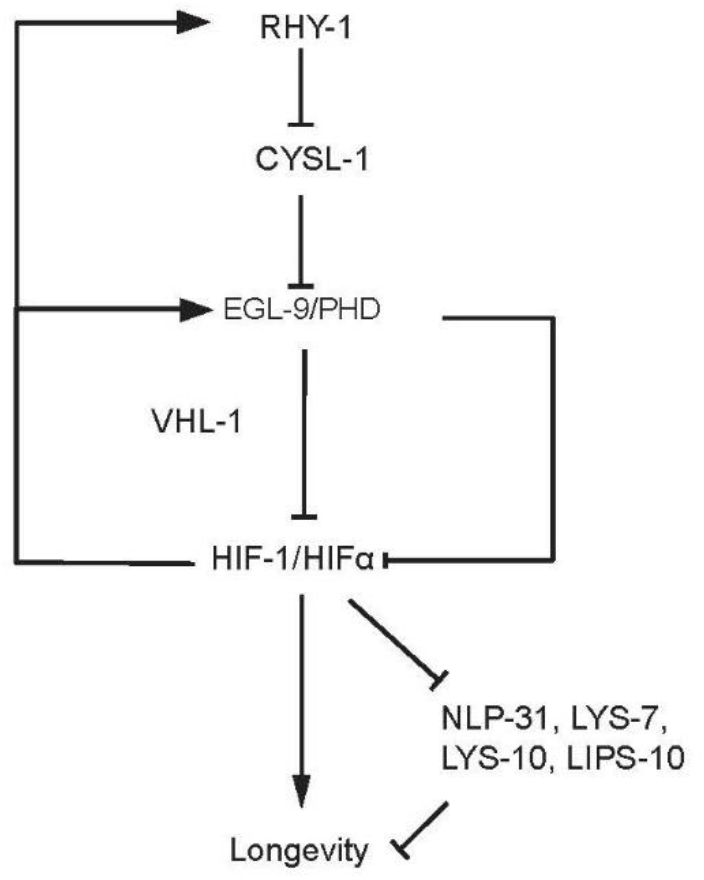
Epistatic model of lifespan regulation by VHL-1, RHY-1, and EGL-9. RHY-1 and EGL-9 act in the same pathway to inhibit HIF-1 activity through both VHL-1-dependent and VHL-1-independent mechanisms. When HIF-1 is stabilized through loss of VHL-1, expression of RHY-1 and EGL-9 is increased, driving reductions in expression of longevity reducing target genes including, NLP-31, LYS-7, LYS-10 and LIPS-10 through a HIF-dependent and VHL-1-independent mechanism. Inhibition of EGL-9 activity causes upregulation of genes including NLP-31, LYS-7, LYS-10 and LIPS-10, inhibiting longevity.

EGL-9 and VHL-1 have distinct roles in pathogen resistance, with *egl-9(sa307)* mutants exhibiting HIF-1-dependent resistance to fast killing by *pseudomonas aeruginosa* while *vhl-1(ok161)* mutants do not (Luhachack et al., 2012; Shao et al., 2010). We noticed that our list of EGL-9/VHL-1 antagonistic genes contained several genes with reported or predicted roles in pathogen response, including *nlp-31*, which encodes five neuropeptide-like proteins with functions in defense against fungal pathogens and gram-negative pathogenic bacteria, *lys-7*, which encodes a lysozyme with a reported role in defense against gram-negative bacteria, *lys-10*, which encodes another lysozyme, and *lips-10*, which encodes an enzyme that, like lysozymes, has hydrolase activity (Couillault et al., 2004; Harris et al., 2010; Marsh et al., 2011; Nathoo et al., 2001). We found that treatment with *lys-10(RNAi), lips-10(RNAi)*, *lys-7(RNAi)*, and *nlp-31(RNAi)* each extended longevity of *egl-9;vhl-1* mutants **(Fig. 4 B,D,F,H)**.

We also tested treatment with RNAi targeting the ferritin genes, *ftn-1* and *ftn-2*, predicted oxidative and heavy metal response genes that have been identified as EGL-9/VHL-1 antagonistic genes in multiple studies (Ackerman & Gems, 2012; Romero-Afrima et al., 2020). We found slight but significant increases in lifespan in *egl-9(sa307);vhl-1(ok161)* animals treated with *ftn-1(RNAi)* and *ftn-2(RNAi)*, however the magnitude of these changes was small and they were not consistent across individual replicates (significant in 2 of 4 trials) (**Figure S2** and **Table S2**). These data are consistent with ferritins playing a minor role in modulation of lifespan by the hypoxic response.

Taken together, these results are consistent with a model in which EGL-9 activity promotes longevity during HIF-1 stabilization through VHL-1-independent inhibition of target genes, including multiple genes with predicted functions in defense against pathogens.

## Discussion

Collectively, our results show that, while reduction in RHY-1 and EGL-9 activity can increase lifespan via the hypoxic response, RHY-1 and EGL-9 activity also promote longevity downstream of HIF-1 stabilization by *vhl-1* mutation. We further demonstrate that RHY-1 and EGL-9 activity are required to control the direction of differential expression of numerous transcripts in *vhl-1* mutants and that EGL-9-dependent suppression of *nlp-31*, *lys-7*, *lys-10*, and *lips-10* promotes longevity in *vhl-1* mutants.

### Genetic interactions of *vhl-1*, *egl-9*, and *rhy-1*

While HIF has emerged as a key regulator of longevity, it was previously unknown how hypoxic signaling pathway components interact to influence longevity. HIF-1 hydroxylation is required for its interaction with VHL-1, so in theory we might expect *egl-9* mutant phenotypes to be epistatic to *vhl-1* mutant phenotypes. However, reports that 1) EGL-9 and RHY-1 have VHL-1-independent roles in the expression and tissue distribution of hypoxic response genes, 2) EGL-9 interacts with other proteins through proline-hydroxylase-activity-dependent and -independent mechanisms to influence phenotype, and 3) EGL-9 and RHY-1 are transcriptionally upregulated when HIF-1 is stabilized, make these phenotypic interactions difficult to predict (Luhachack et al., 2012; Shao et al., 2009; Shao et al., 2010; Shen et al., 2005; Shen et al., 2006).

We found that *rhy-1(ok1402)* blocked lifespan extension by *vhl-1(RNAi)* and *vhl-1(ok161)*, while *egl-9(sa307)* blocked lifespan extension by *vhl-1(RNAi)* and partially blocked lifespan extension by *vhl-1(ok161)*. While these results are broadly consistent with a model where RHY-1 regulates lifespan through its known EGL-9 modulating activity, it is interesting that *rhy-1* mutation has a stronger effect on the longevity of *vhl-1(ok161)* mutants than the *egl-9(sa307)* mutation. This could be explained by the reportedly more robust HIF-1 stabilization and upregulation of pro-longevity HIF-1 target genes in *egl-9(sa307)* relative to *rhy-1(ok1402)*, or by an additional, HIF-1-independent role for RHY-1 in longevity determination (Shen et al., 2006).

We confirmed the surprising, previously reported, result that the *rhy-1(ok1402)* mutation does not extend lifespan, despite stabilizing HIF-1. However, interestingly, *rhy-1(RNAi)* extends lifespan through a mechanism that is partially dependent on its established interactions with CYSL-1 and HIF-1. While off-target effects of RNAi are a concern when mutant and RNAi phenotypes differ, the observation that the *rhy-1(RNAi)*-mediated lifespan increase is fully abrogated in a *rhy-1(ok1402)* mutant background strongly suggests that modulation of RHY-1 is the primary factor influencing lifespan in this context. It is worth noting that, while *cysl-1(ok762)* and *hif-1(ia4)* mutations reduce the longevity promoting effect of *rhy-1(RNAi)*, neither completely abrogates it. This suggests that RHY-1 may have secondary, HIF-1-independent roles that influence longevity. Published reports showing that *rhy-1;hif-1* compound mutants have a synthetic deleterious effect on fertility and that RHY-1 modulates hydrogen-sulfide resistance in a HIF-1-independent manner also suggest interesting HIF-1-independent roles for RHY-1 (Horsman et al., 2019; Shen et al., 2006).

A published mechanism explaining the interaction between RHY-1 and HIF-1 suggests that *rhy-1* mutants should largely phenocopy *egl-9* mutants (Ma et al., 2012). While interpretation of *egl-9* phenotypes is complicated by the lack of a viable *egl-9* null mutant, published results do suggest that *egl-9(n571*), a point mutant that is predicted to affect splicing, and *egl-9(RNAi)* have more robust longevity promoting phenotypes than the strong loss-of-function mutant, *egl-9(sa307)*, a deletion in the EGL-9 catalytic domain (Darby et al., 1999; Miller et al., 2017; Shao et al., 2009; Trent et al., 1983). These results are consistent with EGL-9 also having longevity promoting and limiting functions, with the mutant phenotype depending on the severity of the loss in activity. These data are consistent with a model in which EGl-9 and RHY-1 act in the same pathway to both limit wild-type longevity by destabilizing HIF-1 and increase longevity when HIF-1 is stabilized.

### VHL-1-independent EGL-9 targets modulate lifespan

Along with other groups, we identified a substantial subset of targets that are transcriptionally regulated in opposite directions by EGL-9 and VHL-1 (Ackerman & Gems, 2012; Angeles-Albores et al., 2018). This suggests that upregulation of EGL-9 and RHY-1 when HIF-1 is stabilized has substantial effects on the hypoxic transcriptome in addition to possible feedback regulation of HIF-1. Previous publications have established that EGL-9 has VHL-1-independent activities that can increase or reduce resistance to various pathogens (Luhachack et al., 2012; Shao et al., 2010). Here, we find that several RNAi clones targeting genes with reported or likely functions in innate immunity, *nlp-31(RNAi) lys-7(RNAi)*, *lys-10(RNAi)*, and *lips-10(RNAi)*, extend lifespan in *egl-9(sa307);vhl-1(ok161)* mutants, suggesting that EGL-9-dependent downregulation of these genes promotes longevity in *vhl-1(ok161)* mutants.

The mechanisms underlying VHL-1-independent transcriptional regulation by EGL-9 have not been fully established. Previous studies report that *egl-9(sa307)* and *hif-1(ia4)* loss of function mutants cause transcriptional upregulation of the direct HIF-1 transcriptional target *ftn-1*, while *vhl-1* loss of function mutants and overexpression of non-hydroxylatable HIF-1 (P621A) suppress *ftn-1* expression (Ackerman & Gems, 2012; Romero-Afrima et al., 2020). These results are consistent with a model in which binding of hydroxylated and non-hydroxylated HIF-1 may have opposite effects on the *ftn-1* promoter region, with *vhl-1(ok161)* mutation causing increases in hydroxylated HIF-1 while *hif-1(ia4)* and *egl-9(sa307)* mutation both eliminate hydroxylated HIF-1 (Ackerman & Gems, 2012; Romero-Afrima et al., 2020). A recent analysis from the Sternberg group showed expression patterns consistent with a role for hydroxylated HIF-1 in expression of a larger set of transcripts that are oppositely affected by VHL-1 and EGL-9 (Angeles-Albores et al., 2018).

Other reports suggest that EGL-9 may trigger VHL-1-independent transcriptional responses through more complex mechanisms. Multiple labs report that EGL-9 affects gene expression through mechanisms that are independent of its hydroxylation activity (Luhachack et al., 2012; Shao et al., 2009). One study suggests that EGL-9 represses HIF-1 activity through a mechanism that requires physical interaction between EGL-9/PHD and the WD repeat containing protein SWAN-1 (Shao et al., 2010). EGL-9 might also affect gene expression via hydroxylation of substrates other than HIF-1, with one study indicating that LIN-10 is a target of EGL-9 hydroxylation (Park et al., 2012). Mechanistic biochemical studies to 1) identify EGL-9 hydroxylation targets and protein-protein interaction partners, and 2) establish whether hydroxylated and non-hydroxylated HIF-1 interact with distinct transcriptional complexes, will be needed to fully understand the complex biological function of EGL-9/PHD.

Constitutive sterile activation of the innate immune response increases during mammalian aging and is a key driver of many age-related pathologies (Franceschi & Campisi, 2014). A trade-off between constitutive immune activation and longevity has also been established in *Drosophila* (Libert et al., 2006). In *C. elegans*, immune-related signaling genes contribute to the longevity phenotypes of long-lived insulin signaling mutants; however, to our knowledge, this is the first evidence of antagonism between immune activation and non-pathogen exposed survival in *C. elegans* (Murphy et al., 2003). Further mechanistic studies of the connection between pathogen resistance genes and accelerated aging-related phenotypes in tractable model systems may yield insights and interventions that can be translated to mammalian inflammatory aging pathologies.

## Conclusion

Drugs that inhibit PHD activity are currently in clinical trials for anemia, and animal studies have suggested that they may have efficacy in models of neurodegenerative conditions, indicating that modulation of the hypoxic response to treat age-related disorders in humans may be on the horizon (Ashok et al., 2017; Haase, 2017; Li et al., 2018; Mehta et al., 2009). However, the role of HIF-1 activity in modulating aging and age-related pathologies is highly complex. The finding that RHY-1 and EGL-9/PHD modulate aging by a VHL-1-independent mechanism indicates that further studies to illuminate the mechanisms underlying non-canonical roles of proline-hydroxylases in influencing transcription, aging, and disease will be critical to support effective translation of compounds that modulate the hypoxic response.

## Methods

### Worm culture and RNAi

Standard procedures for *C. elegans* strain maintenance and handling were used (Ahringer, 2006; Stiernagle, 2019). Experiments were performed on animals grown on *Escherichia coli* HT115 expressing empty vector or target RNAi from the Ahringer library. Nematode strains N2, *rhy-1(ok1402)*, *egl-9(sa307)*; *vhl-1(ok161)*, *hif-1(ia4)*, *cysl-1(ok762)* and *egl-9(sa307);vhl-1(ok161)* were obtained from the Caenorhabditis Genetics Center. *rhy-1(ok1402)* was outcrossed 5 times to a recently unfrozen CGC N2 strain, and *rhy-1(ok1402);vhl-1(ok161)* was made from outcrossed *rhy-1(ok1402)* using standard methods.

### Lifespans

Lifespans were performed as described previously with minor modifications (Amrit et al., 2014). Animals were cultured on standard RNAi plates (NGM + 4ml 1M IPTG/L) inoculated with empty vector RNAi or the clone being tested, at the temperature of the assay, for at least two generations prior to measuring lifespans. Synchronized populations were transferred to lifespan plates (NGM+100mg Carbenecillin 4ml1M IPTG/l+660ul 150mM FUDR/L) at adulthood. Animals with age-related vulval integrity defects were included, animals that left the plate were not considered (Leiser et al., 2016). Replicates and statistics are included in supplemental table 1.

### RNA isolation, sequencing and analysis

Worm strains were synchronized by treating gravid adult worms with sodium hypochlorite and collecting ~1000 offspring per genetic condition. Once the offspring reached young adulthood, they were collected in M9 buffer and immediately flash-frozen in liquid nitrogen. RNA was extracted following Invitrogen’s TRIZOL RNA extraction protocol. Before library preparation the samples were analyzed on an Agilent 2100 BioAnalyzer. Only samples with an RNA integrity numbers (RIN) equal to or greater than 9.0 were used in downstream analyses. Single-strand reverse transcription, library preparation, and sequencing were performed on an Illumina machine. Read alignments were mapped using Kallisto and differential expression analyses was performed using Sleuth. We used a q-value cutoff of <.05 for downstream comparisons across conditions as it’s adjusted for multiple hypothesis testing (Bray et al., 2016; Pimentel et al., 2017).

## Supporting information

table s1. aggregated lifespan data

table s2. lifespan data for individual experiments

table s3. consistent hypoxic response genes in table 1

table s4. differentially expressed transcript in figure 3A

table s5. transcripts in quadrants II and IV of figure 3C

table s6. combined B values in Figure 4

## Acknowledgements

We thank David Angeles and the Sternberg lab for sharing their Kallisto analyses and expertise. This work was supported by NIH R01AG058717 and the Glenn Foundation for Medical Research. HAM was supported by NIH F31AG060663. Additionally, we thank the University of Washington Nathan Shock Center for support of gene expression analysis (NIH P30AG013280). Strains were provided by the Caenorhabditis Genetics Center that is funded by the NIH ORIP (P40OD010440).

---

**Figure S1.**
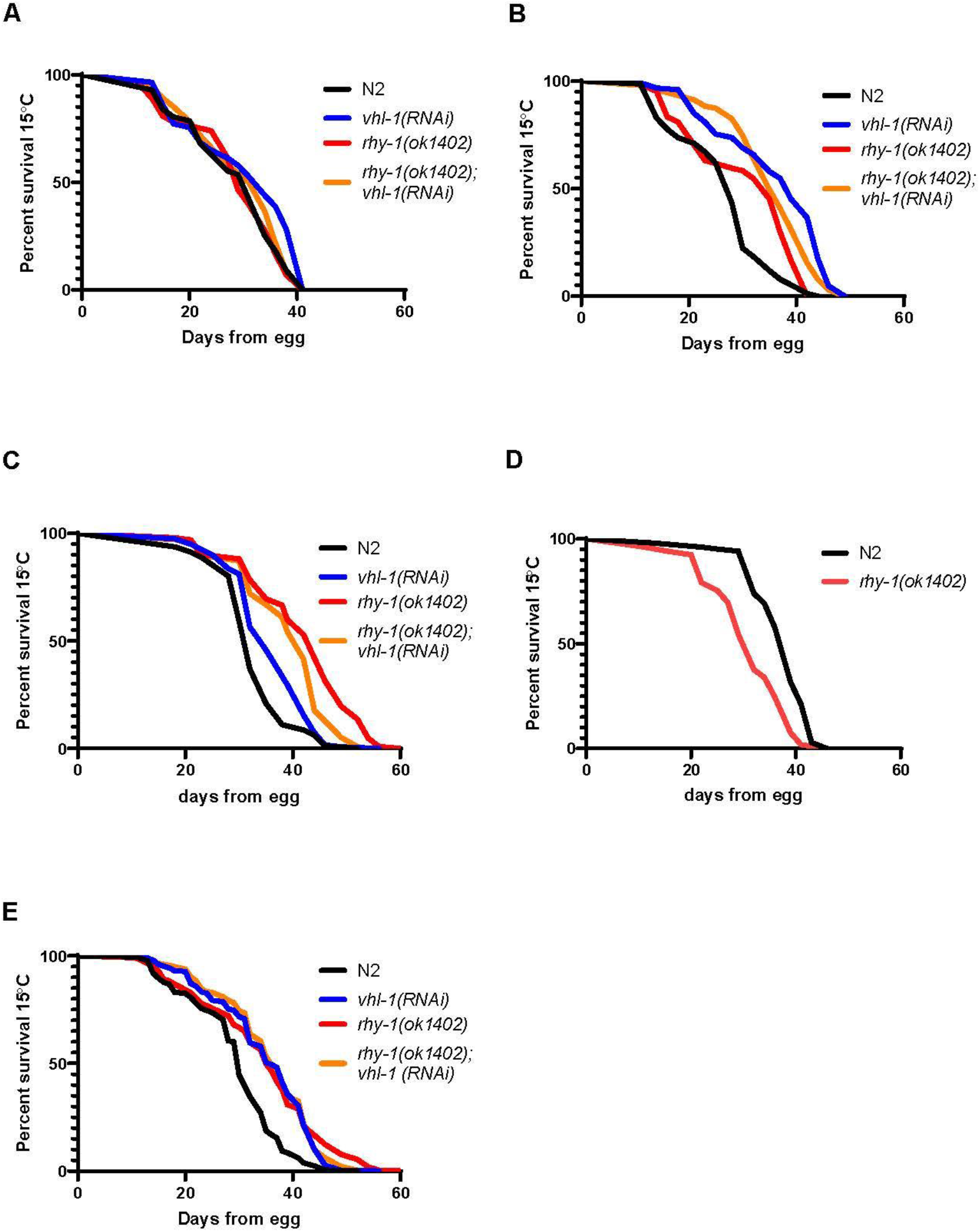
rhy-1-mediated effects of lifespan. A-C) Individual lifespan experiments with N2 and rhy-1(ok1402) animals treated with empty vector or *vhl-1*(*RNAi*). D). Additional lifespan trial comparing N2 and rhy-1(ok1402) lifespans on empty vector RNAi. E. Aggregated data from A-C.

**Figure S2.**
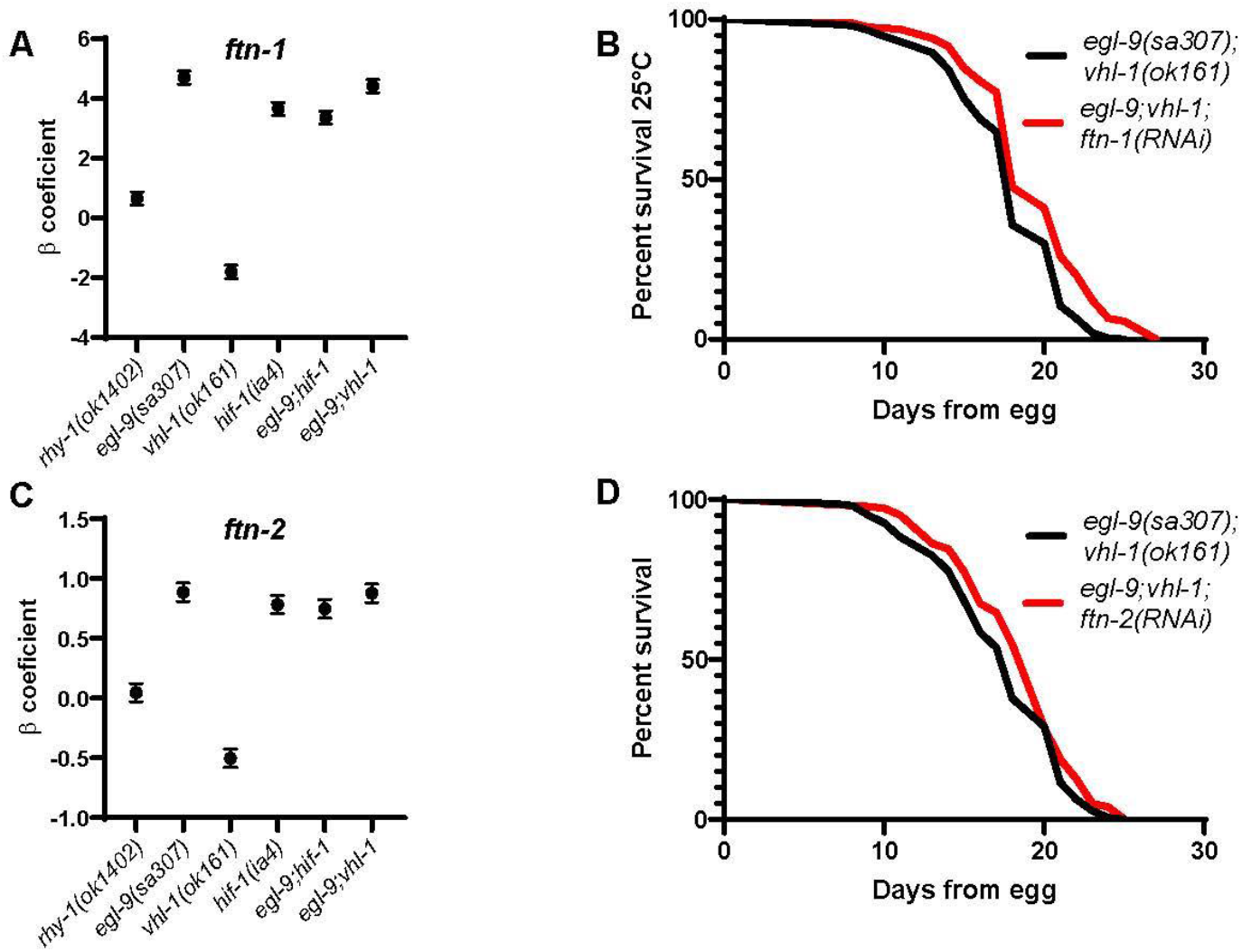
Effects of additional VHL-1/EGL-9 antagonistic HIF-1 targets on lifespan. A, C) expression levels of selected transcripts from RNA-seq analyses. B-D) Treatment with *ftn-1*(*RNAi*), and *ftn-2*(*RNAi*) increases lifespan of *egl-9*(*sa307*);*vhl-1*(*ok161*) mutants (p<.05 by log-rank). Lifespan data are aggregated from at least three experiments, and are significant (p<.05 by log-rank with Bonferroni correction) in 2 out of 4 individual trials (Table S2).

